# Temporal Indexing of Motor Memory

**DOI:** 10.64898/2026.09.10.750443

**Authors:** Apoorva Sharma, Hanna Hillman, Samuel D. McDougle

**Author notes:** Correspondence: Apoorva Sharma.

## Abstract

Humans must store and retrieve distinct implicit motor memories in different contexts. Recent research has revealed the motor adaptation system’s sensitivity to contextual cues related to posture, spatial goals, and visual inputs. However, to date, the role of one critical variable in implicit motor memory retrieval – the passage of time – remains underexplored, even though it is arguably more fundamental. Here we ask if different temporal intervals can act as effective contextual cues for retrieving competing motor adaptation memories at the subsecond level. We combined a visuomotor dual adaptation paradigm with a ‘set-go’ task to link opposite perturbation contexts with different movement preparation intervals. The preparation intervals (and their associated perturbations) were randomly interleaved during training, leading to consistent interference. After training, a testing phase revealed robust contextual modulation effects, such that distinct implicit motor memories were retrieved based on the time elapsed during the pre-movement preparatory interval. These contextual memory effects generalized to novel preparatory intervals in a highly structured fashion. Contextual motor memory separation was seen both when elapsed time had to be internally tracked (high uncertainty) and when it was aided by visual and symbolic cues (low uncertainty). Learning and generalization data across experiments could be parsimoniously explained by a computational model that indexed motor memories via a cerebellar granule-cell-like temporal basis set that tiled the delay interval. Our findings suggest that intrinsic timing processes in the cerebellum may function to index motor memories and generalize those memories to new temporal contexts.

## Introduction

Humans are often required to rapidly and automatically select appropriate motor memories – from deciding the exact pressure to apply to a car brake while approaching a traffic light, to retrieving different baseball swing kinematics when trying to hit a changeup versus a fastball. One important goal in the field of motor control is characterizing the type of contextual cues that help the motor system select among competing motor memories prior to movement. Cues considered in previous research include variables such as posture, the spatial origin or final goal of movements, sensory signals (e.g., light flashes and tones), and even cognitive states like decision uncertainty and attention (Avraham et al., 2022; I. S. Howard et al., 2013; Hwang et al., 2006; Krouchev & Kalaska, 2003; Ogasa et al., 2024; Sheahan et al., 2016; Wang et al., 2025). Such cues often rely on either salient sensory variables or task states (e.g., dynamic visual stimuli) or motoric variables (e.g., postures, lead-in and follow-through movements, etc.).

Despite these advances, the passage of time remains relatively untested as a contextual cue for motor memory. This is an important gap, as dynamics play a central role in the preparation and execution of movements. A popular idea in modern motor neuroscience is that sensorimotor cortex functions as a dynamical system that is optimized to shape time-varying muscle commands (Shenoy et al., 2013). Movement preparation, in turn, is thought to be guided by its own set of cortical dynamics, which primarily function to ‘seed’ the initial state of subsequent movement-related neural dynamics (Churchland & Shenoy, 2024; Heald et al., 2021; Sheahan et al., 2016). Inspired by these ideas, here we investigated whether the length of movement preparatory periods can serve as a robust cue for selecting among distinct competing motor memories. We hypothesized that distinct temporal contexts, by directly influencing motor planning dynamics, may help index competing motor memories, which are thought to be represented as distinct engrams in the cerebellum (Mauk & Ruiz, 1992; Perrett et al., 1993).

We implemented a visuomotor dual adaptation paradigm in which human participants made reaching movements to a visual target and experienced feedback perturbations that were randomly distributed over training to create interference between competing implicit motor memories. Participants were made aware of the perturbed feedback but actively ignored it, ensuring that the resulting adaptation memory was implicit (Huberdeau et al., 2015; O. A. Kim et al., 2022; Morehead et al., 2017; Taylor et al., 2014). Embedded in our adaptation paradigm was a ‘set-go’ task (Jazayeri & Shadlen, 2015), where each perturbation context was deterministically coupled with a distinct preparatory interval – the delay between a ‘set’ and a ‘go’ cue.

To preview our results, we found in post-training tests that people effectively partitioned motor memories across the delay contexts. Moreover, generalization tests showed an orderly pattern of behavior when novel preparatory intervals were introduced, demonstrating that the temporal indexing of motor memory generalizes in a highly structured fashion. In further experiments we used dynamic looming cues and symbolic numeric cues to reduce the uncertainty in the amount of elapsed time during movement preparation. Both methods for reducing uncertainty led to stronger contextual memory separation effects. We performed computational modeling analyses to contrast two competing models of the observed learning and generalization effects: one that linked the effects directly to people’s subjective estimates of elapsed time, and another that framed the effects as intrinsic consequences of hypothesized temporal basis functions manifested in the granule cell layer of the cerebellum (Narain et al., 2018). The cerebellar model outperformed the subjective timing model. Together, our results point to the passage of time during movement preparation – on the order of fractions of a second – as an important feature in motor memory formation and expression.

## Results

### Temporal indexing of motor memory via internal monitoring of elapsed time

We tested whether temporal intervals during movement preparation can serve as an effective context for partitioning competing motor memories. Participants performed a dual-adaptation task in which each planning delay was deterministically paired with a specific error direction. For a given participant, the target appeared only on one side (left or right) and only one delay-direction pairing was used throughout (both target side and delay-direction pairing were counterbalanced across participants).

We first examined behavior during the training block, where both trial-by-trial adaptation and temporal contextual learning were expected to contribute to motor behavior. To dissociate these effects, we fit a linear mixed-effects model predicting participants’ hand angles from the current trial context, one-trial-back rotation direction, two-trials-back rotation direction, trial number, and their interactions, with participant-specific random intercepts and slopes for these effects. As expected, the adaptation effect was significant (β = 2.28, SE = 0.18, *t*(7689) = 12.38, *p* < .001, 95% CI [1.92, 2.64]), demonstrating that the sign of the error on a given trial drove the hand in the opposite direction on the subsequent trial (**Figure 2A**).

Critically, current trial context also significantly modulated hand angles even after accounting for adaptation. **Figure 2B** depicts the context effect across training bins (with adaptation regressed out of hand angle on a per-participant basis, see *Methods*), revealing a gradual increase in contextual memory separation over time (Sheahan et al., 2016). The mixed-effects model revealed a significant overall context effect (β = 1.77, SE = 0.32, *t*(7689) = 5.56, *p* < .001, 95% CI [1.15, 2.40] (**Figure 2C** left panel)) and a significant context X trial number interaction (β = 0.0039°/trial, SE = 0.0013, *t*(7689) = 3.07, *p* = .002, 95% CI [0.0014, 0.0064]) (**Figure 2C** right panel).

Our primary analysis focused on the test phase, where no visual feedback was given and thus further adaptation and context learning were precluded (**Figure 2D-E**). This key phase allows for a clean, direct test of contextual motor memory separation, unconfounded by learning and/or rotation schedule effects. We first focused on the two delays at which the participants were trained, 500 and 1500 ms, reasoning that a reliable difference between them at test would reflect expression of the trained contextual associations. For each participant, we computed the (baseline-subtracted) difference in hand angle between these two delays in the test phase (**Figure 2D**). This difference (after aligning the two counterbalanced subgroups) was significantly different from zero (M = 1.578 ± 2.283°, *t*(21) = 3.242, *p* = .004).

We also asked whether the delay-dependent context effect generalized to novel delays. To remove between-subject offsets in hand angle or biases, we mean-centered each participant’s data across the five delays and then averaged trials within each delay for each participant. We observed a strikingly graded pattern for both counterbalanced subgroups (**Figure 2E** left panel): Subgroup 1 (CW/CCW) showed a transition from positive to negative hand angles with increasing delay and the counterbalanced subgroup 2 (CCW/CW) showed the reciprocal pattern. When the two subgroups were aligned, the overall graded pattern was evident (t-tests against 0 context effect; 500 ms: M = −1.025 ± 1.291°, *t*(21) = −3.722, *p* = .001; 750 ms: M = −0.340 ± 0.851°, *t*(21) = −1.872, *p* = .075; 1000 ms: M = 0.088 ± 0.719°, *t*(21) = 0.575, *p* = .571; 1250 ms: M = 0.771 ± 1.064°, *t*(21) = 3.397, *p* = .003; 1500 ms: M = 0.748 ± 0.857°, *t*(21) = 4.094, *p* = .001) (**Figure 2E** right panel). These results demonstrate that the learned temporal-error associations generalized to new temporal intervals not experienced during training.

### Computational modeling of temporal indexing effects

The graded generalization pattern we observed suggests that participants represent elapsed time as a continuous variable linked to motor memory rather than learning two isolated associations at 500 and 1500 ms. To ask what kind of temporal representation could produce this pattern, we built two computational models that differed in how elapsed time was represented and linked to motor memory. We fit both models to participants’ training phase data and compared their out-of-sample predictions against the observed test phase data.

Prevailing models of temporal anticipation have addressed how an agent estimates a probability distribution of observing sensory events at certain times, and the associated dynamic ‘hazard function’ that reflects the instantaneous rate at which a specific event is expected to occur and which can be used to estimate response latencies (Janssen & Shadlen, 2005; Luce, 1991; A. Nobre et al., 2007). Meanwhile, associative learning models of both trial-by-trial motor adaptation and tasks like eyeblink conditioning have been designed to link specific sensory events with specific memory states, and are well suited to capture contextual tagging of motor memory (Avraham et al., 2022; Mauk & Ruiz, 1992; Rescorla & Wagner, 1972). We combined these two approaches in a single model that explicitly represented delay intervals (hazard rate component) and associated those intervals with different motor memory states (**Figure 2F**).

The subjective timing model functions by assuming that participants learn a subjective probability density function of expected go-cue times during the early phases of the task, with classic Weber-scaled uncertainty in timing (see *Methods* for further details of the computational model). Consistent with this account, participants responded more slowly following the shorter 500 ms preparatory delay than the longer 1500 ms delay (mean participant median RT: 408.64 ms vs 347.57 ms, respectively; *t*(21) = -8.15, *p*< .001). The hazard function is then computed from the subjective PDF, after fitting participants’ training phase RT data to estimate a Weber fraction (fit Weber fraction values: 0.130 ± 0.010).

Motor learning data were modeled using error-based learning, where a delay-agnostic memory state and two delay-specific states (one per trained interval) were each updated by the experienced error, thus capturing both trial-by-trial adaptation and incremental context learning (see *Methods*). This gradual contextual learning was integrated into the modeled timing computations, allowing us to simulate a representation of the agent’s rotation direction prediction as a continuous function of elapsed time, both within trials and across learning. After fitting the model to the training phase data, the model could then be used to simulate behavior in the critical testing phases, by taking the learned continuous-time rotation prediction representation and ‘reading off’ that prediction at both the learned and novel delay intervals probed in the testing phase. Across all five testing delays, the out-of-sample model predictions decently matched the shape of the testing phase data (R^2^ = 0.829) even though it was not fit to those data (**Figure 2H**). The subjective timing model thus captured the key observed behavioral effects in this experiment.

As an alternative account of how elapsed time supports context-dependent learning, we adapted a temporal basis set framework proposed for cerebellar granule cells (Narain et al. 2018). Here, modeled granule cells respond maximally at different elapsed times, together forming a heterogeneous population of Gaussian temporal kernels (**Figure 2G**). Estimates of elapsed time arise from the weighted combination of activity across these temporal kernels, such that learning can generalize smoothly to time points that were never directly experienced during training. We implemented this using a fixed set of overlapping Gaussian basis functions spaced continuously across the delay period, with kernel width increasing linearly for later-centered kernels (see *Methods* for parameters).

As in the subjective timing model, learning included a delay-agnostic state that updated on every trial. The delay-specific component was a vector weight over the temporal basis set, updated by the same error-based rule, so that learning on a given trial adapted only the weights of basis elements active near that trial’s delay. Predicted hand angle was the sum of delay-agnostic state and the weighted basis activity at that trial’s delay.

Since this basis spans the full range of delays continuously, generalization to novel test-phase delays followed directly from evaluating the learned weights at any elapsed time **(Figure 2G**). Across all five testing delays, these out-of-sample predictions matched the shape of the test phase data (R^2^ = 0.969). Model prediction error across the untrained delays was comparable for the granule-cell basis model and the subjective timing model (mean RMSE = 0.830 vs 0.831, respectively; Wilcoxon signed-rank test, *W* = 118, *p* = .627). Fitted parameter values for these modeling results are presented in **Figure S1**. Overall, these findings establish two ways of thinking about temporal indexing in motor adaptation – either as a function of a subjective timing process, or as an intrinsic and obligatory consequence of a temporal basis set (i.e., in the cerebellum) that automatically links elapsed time to different motor memories. Our next experiment was designed to distinguish which model explained our data better by reducing the subjective uncertainty in elapsed time.

### Temporal indexing under reduced uncertainty

Building on the robust context separation we observed when time had to be internally-monitored, we next asked if and how reducing temporal uncertainty would affect contextual separation of motor memories. We achieved this by explicitly cueing the upcoming delay using a looming target: the target grew in size over the course of the delay period, expanding slowly for long delays and rapidly for short delays (**Figure 3A**). Critically, our two models made distinct predictions under this form of reduced uncertainty. The subjective timing model predicted sharply reduced generalization, making the competing motor memories more strongly linked to the exact predicted intervals. Indeed, participants’ RTs were not different across the two delays, leading to a Weber fraction approaching 0 (and thus leading to extremely narrow subjective time PDFs and minimal generalization effects; see below, and *Methods*). The granule-cell model, however, still predicted orderly generalization effects.

As before, we fit a linear mixed-effects model to training block hand angle, with current-trial context, one-trial-back rotation direction, two-trials-back rotation direction, trial number, and their interactions as fixed effects, and random slopes and intercepts by participant. Adaptation again emerged as a strong predictor of hand angle (β = 1.64, SE = 0.19, *t*(8367) = 8.48, *p* < .001, 95% CI [1.26, 2.02]) (**Figure 3B**). After accounting for adaptation, context significantly modulated hand angle (β = 2.58, SE = 0.43, *t*(8367) = 6.01, *p* < .001, 95% CI [1.74, 3.43]) (**Figure 3C-D**) and the context effect grew over the course of training (context X trial number interaction: β = 0.0030°/trial, SE = 0.0010, *t*(8367) = 2.88, *p* = .004, 95% CI [0.0010, 0.0050]).

As in experiment 1, our primary analyses focused on the test phase. We first compared the hand angles at the two trained delays, 500 and 1500 ms (**Figure 3E**). The baseline-subtracted difference between these delays was significantly different from zero (M = 3.699 ± 3.543°, *t*(21) = 4.896, *p* < .001). The graded pattern observed previously was evident again under reduced temporal uncertainty (**Figure 3F**, left panel). Analysis of the aligned generalization functions confirmed a graded pattern (500 ms: M = −1.854 ± 1.442°, *t*(21) = −6.031, *p* < .001; 750 ms: M = −0.541 ± 0.632°, *t*(21) = −4.010, *p* = .001; 1000 ms: M = 0.637 ± 0.503°, *t*(21) = 5.939, *p* < .001; 1250 ms: M = 0.847 ± 0.849°, *t*(21) = 4.678, *p* < .001; 1500 ms: M = 1.026 ± 1.144°, *t*(21) = 4.206, *p* < .001) (**Figure 3F** right panel).

As in the previous experiment, we compared the two models in their ability to predict the held out test phase data. Since participants in this experiment received an explicit cue indicating the upcoming delay, the temporal uncertainty that the subjective timing model is built to capture was effectively eliminated, expressed by nearly equivalent RTs across the intervals (mean participant median RT: 275.00 ms at 500 ms vs 286.02 ms at 1500 ms; t(21) = 1.98, p = .061). Because the RT pattern was flattened relative to a standard interval-timing task that does not have explicit timing cues, we fixed the Weber fraction to a near zero value (0.001), reflecting negligible timing uncertainty. As a result, the model’s test phase predictions show no generalization to the novel delay intervals and fit the average data only moderately well, as the model still predicts context effects at the learned intervals (**Figure 3G**; R^2^ = 0.692).

In contrast, the granule-cell basis model’s out-of-sample predictions closely matched the observed generalization pattern (R^2^ = 0.943). Participant-level comparison between the two models’ prediction errors at the untrained delays revealed a significantly lower RMSE for the granule-cell basis model versus the subjective timing model (mean RMSE: 0.633 vs 0.856, respectively; Wilcoxon signed-rank test: *W* = 197, *p* = .022). Fitted parameter values for these modeling results are presented in **Figure S2**. The results of this experiment demonstrate that reducing temporal uncertainty increases the strength of contextual memory separation in our task. Moreover, our observed generalization effects could not be explained as a function of subjective timing processes, but rather appear to be an obligatory consequence of intrinsic timing representations.

### Temporal uncertainty reduction via symbolic cueing

The previous experiment used a dynamic perceptual cue to decrease participants’ uncertainty about elapsed time. This result raised an additional question – does the effect of temporal uncertainty reduction on improving memory separation depend on dynamic perceptual input rather than the passage of time? That is, can uncertainty reduction via explicit, symbolic cues (instead of dynamic perceptual cues) also boost memory separation relative to the high-uncertainty internal timekeeping in our first experiment?

In a follow-up study, we replaced the looming cue with a discrete numeric cue to test whether enhanced temporal indexing persists when uncertainty is reduced but an internal timekeeping process is still required (**Figure 3H**). Participants were instructed to attend to the numeric context cue presented within the reach target, and were informed that the magnitude of the number cue presented was predictive of the delay interval, with higher numbers corresponding to longer intervals.

In the key test phase, comparing the baseline-subtracted test phase hand angles at 500 versus 1500 ms revealed a robust context effect (M = 4.069 ± 3.346°, t(8) = 3.648, *p* = .007) (**Figure 3I**). Additional results of this follow-up study, which include generalization to unseen cues and delay intervals, are depicted in **Figure S3**. Overall, the test phase context effect closely matched the results of the looming-cue experiment, suggesting that internally- and externally-guided timing processes can similarly shape the temporal indexing of motor memories.

## Discussion

The capacity to adaptively retrieve memories in a context-sensitive manner is a fundamental aspect of cognition. The motor system’s similar ability to leverage contextual cues for adaptive memory retrieval has also been documented, primarily for spatial, postural, and sensory contextual cues (Avraham et al., 2022; I. S. Howard et al., 2013; Sheahan et al., 2016). Our findings significantly build on these findings by revealing that the passage of time itself can serve as an effective contextual signal for partitioning motor memories, in a manner consistent with theorized computational functions of the cerebellum.

Across experiments, we examined whether the length of a movement preparation interval could function as a contextual signal for partitioning motor memories, and how this process might be shaped by temporal uncertainty. Under high temporal uncertainty (i.e., internally monitored time with no cues), participants learned to associate distinct preparatory delays with opposing errors (**Figure 1**), producing robust context-dependent motor memory partitioning that persisted into a critical test phase (**Figure 2**). Two additional experiments using either a continuous looming target or discrete numeric cues to signal temporal context (**Figure 3**) also showed robust motor memory separation effects, and increased the size of these effects by nearly 2-fold relative to the high uncertainty condition. All experiments included untrained delays during testing, allowing us to characterize how temporally-associated motor memories generalize to new temporal intervals. Motor behavior at these untrained delays showed graded generalization patterns (**Figure 2, 3**), echoing the parametric spatial generalization observed when motor memories are probed at nearby movement directions (McDougle et al., 2017; Poggio & Bizzi, 2004).

**Figure 1:**
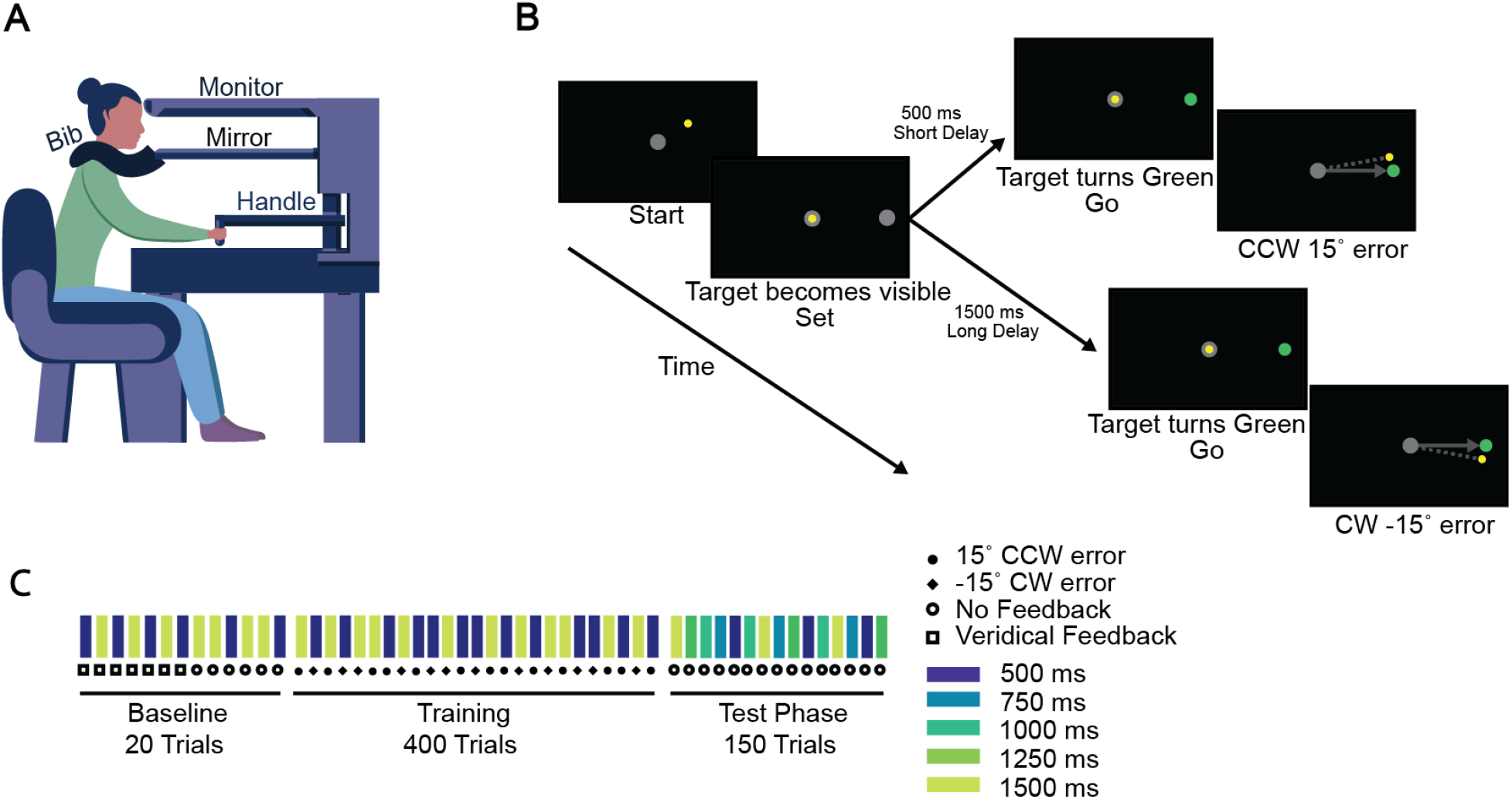
Task design. (**A**) Schematic of the Kinarm robot setup. A horizontal opaque screen blocked any direct view of the participant’s arm. A bib around the neck further blocked any direct view of the body. The task is displayed on the screen and the participant controls the manipulandum below with their right hand. (**B**) The ‘set-go’ task: After the robot centered the hand each trial, a grey target appeared in a fixed position. The appearance of this target (the ‘set’ cue) acted as the start of the planning delay (‘foreperiod’), with the target changing color from grey to green acting as an imperative for participants to move to the target as fast as possible (the ‘go’ cue). The temporal delay between the set and go cues was either 500 ms or 1500 ms during the baseline and training blocks. Each delay was deterministically paired with a specific visuomotor error direction (CW or CCW), where the cursor moved in a fixed (‘clamped’) trajectory, matched in velocity but +/-15° offset from the reach direction. Participants ignored this feedback and planned their reaches directly to the fixed target on all trials, allowing us to measure implicit adaptation effects (see *Methods*). (**C**) Trial schedule: the task started with a baseline block of 20 trials, followed by a training block of 400 trials, and finally the critical test phase (150 trials). No feedback was given in the test phase, and three additional unseen temporal intervals (750, 1000, and 1250 ms) were introduced to measure generalization effects.

**Figure 2:**
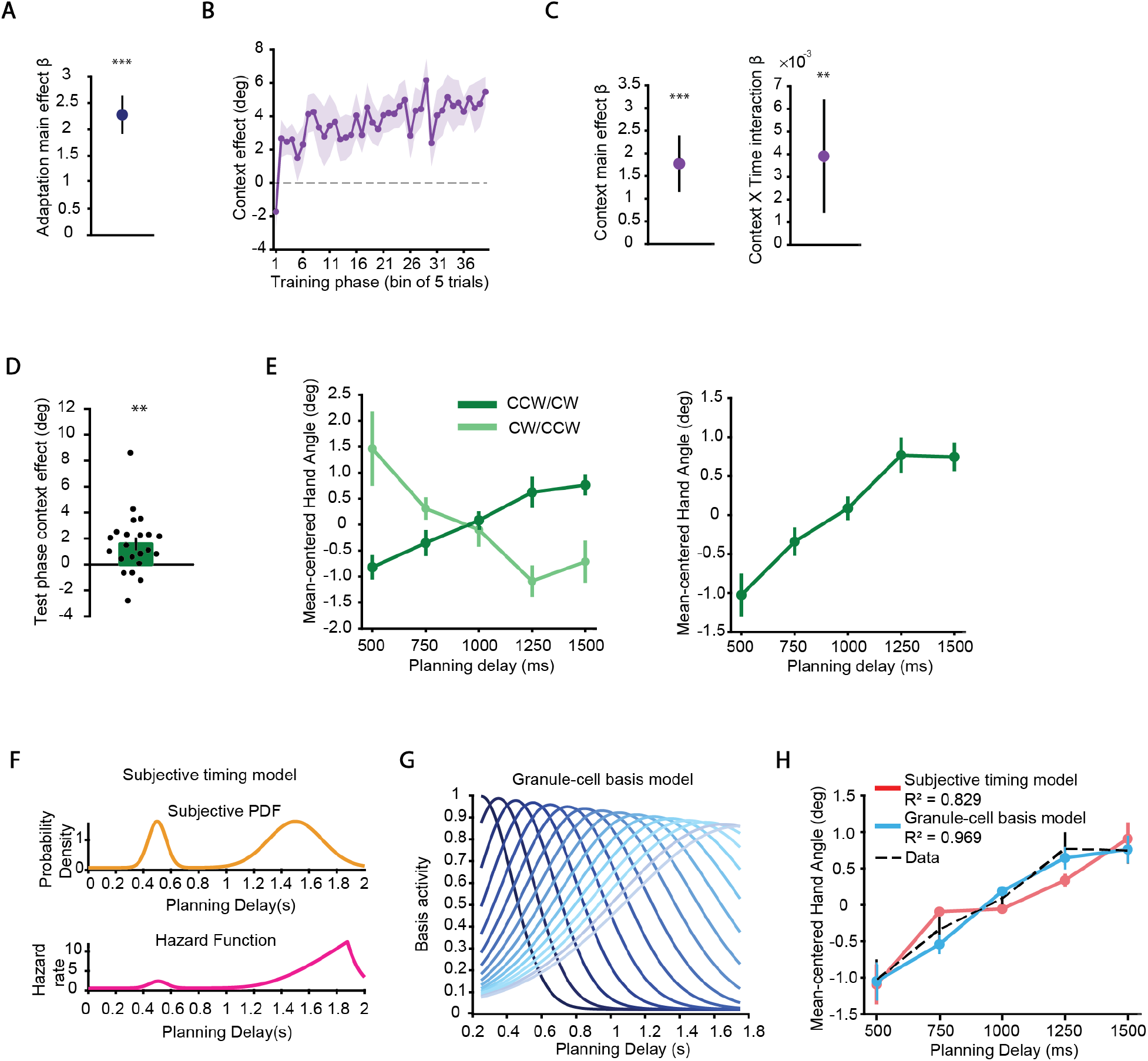
Results: Temporal Indexing under high uncertainty. (**A**) Adaptation main effect β coefficient (from LME model analysis) in the training phase, reflecting the effect of the previous trial’s error direction on the current trial’s hand angle. The positive value reflects compensatory hand angle changes. (**B**) Context-dependent hand angle separation across training bins (i.e., binned differences in hand angle between the two delay conditions), after accounting for adaptation effects. (**C**) Context effect coefficients during training. Left: context main effect β coefficient. Right: context X trial # interaction β coefficient, reflecting growth of the context effect across the training phase. **(D)** Context-dependent hand angle separation (500 ms versus 1500 ms trials) in the test phase. Bar graph depicts the group mean context effect, with individual points showing each participant’s effect. (**E**) Test phase generalization. Left: mean-centered hand angles across the five test delays, with the two counterbalanced delay-rotation mapping subgroups plotted separately. Right: the same test phase data after aligning the two mapping subgroups. **(F)** Subjective timing model: Subjective probability density over expected go-cue times and the corresponding hazard function used to represent internally monitored elapsed time (see *Methods* for model details). (**G**) Granule-cell temporal basis model: Overlapping temporal basis functions (with progressively broader tuning for longer intervals), providing a delay-line representation of elapsed time. (**H**) Model predictions of test-phase generalization data (after being fit on the training phase data). Error bars = 1 S.E.M. \*\**p* < .01, \*\*\**p <* .001

**Figure 3:**
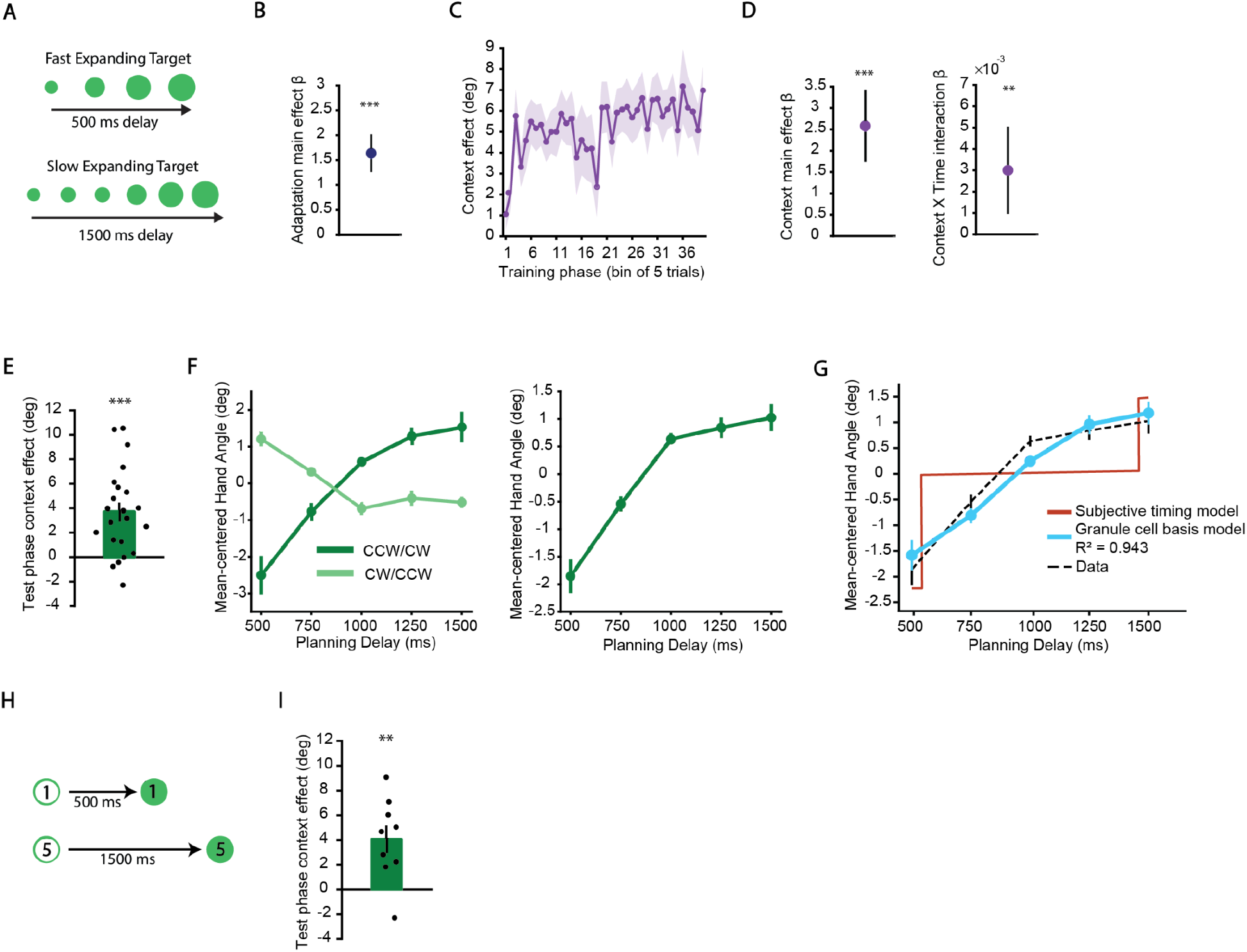
Results: Temporal Indexing under reduced uncertainty. (**A**) Looming cue experiment schematic. The target expanded in size at different rates to explicitly signal the upcoming planning delay (500 ms delay results in faster expansion, 1500 ms delay results in slower expansion of the target). (**B**) Adaptation main-effect β coefficient from the training phase. (**C**) Context-dependent hand angle separation across training bins, after accounting for adaptation effects. (**D**) Regression estimates of contextual effects during training. Left: context main-effect β coefficient. Right: context X trial # interaction β coefficient. (**E**) Context-dependent hand angle separation in test phase. Bar depicts the group mean difference and individual points show each participant’s effect. (**F**) Test-phase temporal generalization. Left: mean-centered hand angle across test delays for the two counterbalanced subgroups plotted separately. Right: same test-phase data after aligning the two subgroups. (**G**) Model predictions in the test phase. (**H**) Discrete numeric cueing experiment schematic. Number cues presented within the reach target explicitly indicated the upcoming planning delay. (**I**) Context-dependent hand angle separation in the test phase. Bar graph depicts the group mean context effect, with individual points showing each participant’s effect. Error bars = 1 S.E.M. \*\**p* < .01, \*\*\**p <* .001

To formalize hypotheses about the timing and memory mechanisms that drove our results, we compared two computational models: one based on established theories about the subjective timing process itself, in which temporal delays modulate motor memory expression and generalization through a hazard-function mechanism that is sensitive to estimated temporal uncertainty (Janssen & Shadlen, 2005; A. Nobre et al., 2007), and another in which temporal context is linked to motor memories via a population of granule-cell-like temporal basis functions (Narain et al., 2018). The granule-cell model best captured behavior across experiments, suggesting that subjective timing computations, while critical for generating temporal predictions and performing explicit timing tasks, are not the key factor driving the temporal indexing of implicit motor memories; rather, motor memory indexing effects are likely an intrinsic and obligatory consequence of a temporal basis-like representation, putatively in the cerebellum.

Is time just another variable that can act as a context for motor memory, or is it special? Previous work has shown that competing motor memories can be indexed by changes in posture, movement direction, spatial goals, sensory cues, or preceding and follow-through movements (Avraham et al., 2022; I.S. Howard et al., 2013; Sheahan et al., 2016). These variables typically alter some aspect of the movement being prepared or the sensorimotor state in which it is prepared. Elapsed time is, of course, more fundamental, continuously evolving even when the planned movement remains unchanged. If motor memories can be indexed simply by when a movement occurs, as our results suggest, this would imply that the motor system can use an internally generated estimate of temporal context to distinguish otherwise identical sensorimotor states. Emerging results in force-field adaptation also point in a similar direction, showing that motor memories can be organized by a movement’s relative (and absolute, as in our study here) progress within a planned action rather than by limb state alone (Makino et al., 2026). In our view, time thus provides a particularly revealing test case of how flexibly the motor system can partition experience into distinct memories that go beyond observable features of the movement and environment.

Our results bridge an important gap between motor learning and other domains of cognition where temporal information has been recognized as an effective contextual cue. For example, in episodic memory research, time has been established as a primary organizing dimension: The hippocampal system employs specialized time cells that fire at specific moments during temporally structured experiences, effectively timestamping memories and distinguishing otherwise similar events (Buzsáki, 2013; Eichenbaum, 2014). These cells create a temporal context that helps organize memories along a continuum, enabling the retrieval of experiences that occurred in close temporal proximity (M. W. Howard & Kahana, 2002). Similarly, research on ‘temporal attention’ has established that predictable temporal structures enhance perceptual processing at anticipated moments in time and even specific locations in space (Correa et al., 2004; Coull & Nobre, 1998; A. C. Nobre & van Ede, 2018; Olson & Chun, 2001). Other work has extended these ideas into the motor domain, showing that implicit temporal expectations can elicit effector-specific motor preparation before an imperative stimulus occurs (van Elswijk et al., 2007; Volberg & Thomaschke, 2017). These findings support the observation that the passage of time can tune motor cortical activity in anticipation of likely actions (Riehle et al., 1997), reinforcing the idea that temporal structure shapes preparatory neural states.

We believe that our results are consistent with a neural mechanism whereby distinct preparatory delays induce divergent trajectories in sensorimotor cortical activity, effectively using time to structure the preparatory state space (Sun et al., 2022). The dynamics of neural population activity are a critical feature in the population code both during movement, and, more relevant to the current study, during movement preparation (Kaufman et al., 2014; Riehle et al., 1997). For example, the exact trajectory that a motor cortical population takes through neural state space during movement depends on its initial state, which is arrived at during preparation. Consistent with this idea, neural activity during movement preparation varies with a wide range of observable kinematic parameters (e.g., Wise et al., 1986; Churchland et al., 2006), even though it does not appear to directly encode those variables (Shenoy et al., 2013).

Recent work has further shown that expectations about upcoming sensory events can systematically shape preparatory population states in M1 and PMd, with trial-to-trial variations in these states predicting subsequent motor behavior (Michaels et al., 2025). What’s more, work in rodents has shown that distinct preparatory neural trajectories can act as contextual indices for similar movement plans (J.-H. Kim et al., 2025).

Given this previous work, we propose that the passage of time during preparation may drive divergent preparatory trajectories in the sensorimotor cortex, allowing different preparatory intervals to act as contextual indices for otherwise similar movements. This timekeeping process may alter the final initial condition for the downstream dynamical system that causes movement generation, and may simultaneously serve as input to the cerebellum, which is a putative key site of the implicit motor memories of interest (Mauk & Ruiz, 1992; Morehead et al., 2017; Perrett et al., 1993) and a crucial neural substrate of subsecond timing computations (Ivry & Keele, 1989; Johansson et al., 2016; Smith, 1968; Trach et al., 2026).

Overall, our findings suggest that the unfolding dynamic of movement preparation can provide structure for separating implicit motor memories. These results raise a number of new questions about the limits and flexibility of temporal context-dependent learning. For example, future work could further manipulate the number of contexts, temporal spacing between contexts, and timing salience to better define the boundary conditions for temporal context-dependent motor learning. Of particular interest is whether an upper limit exists beyond which additional context learning becomes counterproductive. That is, too many contexts compressed within a narrow range may exceed the system’s capacity (e.g., taxing ‘motor working memory’; (McDougle & Hillman, 2025)), resulting in contexts blending together, and perhaps requiring more extensive training to form stable associations. Moreover, engendering different temporal priors in advance of learning by altering the distribution of preparatory timing intervals could help reveal the role of contextual inference processes in the memory indexing process (Heald et al., 2021), and further connect our framework to emerging work on cerebellar associative learning of temporal statistics (Narain et al., 2018).

## Methods

### Participants and Set-Up

Participants (N = 53 after exclusion; mean age = 22.97 years, self reported gender - 31 female, 19 male, 3 prefer not to say) were recruited via Yale’s student psychology pool and received course credit; no financial incentives were provided. All procedures were approved by the Yale University Institutional Review Board, and written informed consent was obtained from each participant prior to participation. Handedness was assessed using the Edinburgh Handedness Inventory (Oldfield, 1971), and no participants were excluded on the basis of handedness.

Participants were seated in front of a KINARM End-Point robotic manipulandum (Ontario, Canada), a low-friction, two-joint device equipped with a cylindrical handle for participants to grasp (**Figure 1A**). The chair was positioned at the center of the manipulandum to ensure that the participant’s body midline aligned with the center of the robotic device. The chair was then adjusted to allow participants to comfortably hold and move the handle, and its height was modified to ensure they could rest their forehead on a head support affixed to the display monitor. Once the desired position was achieved, the chair was locked in place to maintain stability throughout the experiment.

The task was displayed on a screen parallel to the movement plane, reflected from the inverted monitor mounted above. A bib was used to occlude participants’ view of their arms, ensuring that the only visual feedback they received was from the display screen. Participants held the manipulandum’s handle with their right hand in a firm grip, and were instructed not to loosen their hold or release the handle during the experiment. Their left hand rested on a support surface to minimize movement. The entire experiment was conducted in the presence of the experimenter to ensure participants maintained a consistent posture and adhered to the instructions. If a participant loosened their grip on the handle, released the handle, or changed their posture significantly, they were promptly corrected to ensure reliable data collection.

### Task Design

The task workspace consisted of a start position (0.5 cm radius, light grey), a target circle (0.5 cm radius, grey), and a cursor (0.15 cm radius, white) for feedback when required. These elements were displayed on the mirrored screen that participants viewed during the task. Participants made ballistic reaching movements from the start position to the target (distance = 9 cm) while holding the handle of the manipulandum (**Figure 1B)**. Across all experiments, participants were instructed to initiate a rapid reach toward the target at the go cue and were told that precisely stopping at the target was not necessary; they could overshoot the target as long as the movement was completed in time (<500 ms). Each participant only reached to a single target location throughout the experiment. The target could either be positioned directly to the right of the central start position (0°) or the left (180°). Target location and the assignment of 500/1500 ms delays to CW/CCW rotation signs were counterbalanced across participants.

The nature of the go-cue differed across three experimental designs of a “set-go” task. In the internal-monitoring experiment, the target initially appeared grey (the set phase) and changed to green after either a 500 or 1500 ms planning delay, with the color change serving as the go-cue. Thus, participants had to internally monitor the elapsed time during the delay to successfully anticipate the go-cue. In the looming-cue experiment, the target gradually expanded during the planning period, with its rate of expansion precisely indicating the upcoming delay. Participants were instructed to initiate the reach once the target reached its full size (indicated by a thin grey outline marking the target’s final diameter). Finally, in the discrete-cueing control experiment, a number presented within the target symbolically indicated the upcoming wait time. Numbers 1 through 5 corresponded to the five possible planning delays (500, 750, 1000, 1250 and 1500 ms), and participants were informed of this correspondence beforehand. Apart from the above differences in the delay and go-cue features, all experiments were identical.

All experiments consisted of three distinct phases: baseline, training, and test (**Figure 1C)**. Each phase began with a brief pause, during which participants were shown instructions on the screen. In the baseline phase, participants completed 20 trials. For the first 10 trials, they received veridical feedback, with the cursor accurately representing their hand movement throughout the reach (online feedback). The remaining 10 trials were no-feedback trials, requiring participants to reach the target without visual guidance from the cursor.

In the training phase, participants completed 400 trials, and each trial included a fixed cursor rotation of either +15° or -15°. Temporal delays were deterministically coupled with the two rotation directions, and reversed in the second group to counterbalance associations. Participants were instructed to focus on the hand movement from the start position to the target while ignoring the manipulated cursor feedback to ensure implicit learning (Morehead et al., 2017). The same delay-rotation pairing and counterbalancing procedure was used across all three experiments and they differed only in how information about the upcoming delay was provided.

The final test phase in all experiments involved 150 no-feedback trials to test contextual motor memory without the potentially interfering factors of errors or further learning, and to also test generalization to unseen delay intervals. Participants experienced the original short and long delays interleaved with three new delays (750 ms, 1000 ms, and 1250 ms). In all testing phases, each delay occurred equally often and delays were experienced in a random order. Each experiment consisted of 570 trials and took approximately 45 minutes to complete.

### Statistical Analysis

All statistical analyses were performed in MATLAB (R2023a, MathWorks). Two-tailed one-sample t-tests and two-tailed paired *t*-tests were used for planned comparisons between conditions of interest with *α* set to 0.05. Unless otherwise specified, data are reported in the text as mean ± standard deviation (SD); error bars in figures represent standard error of the mean (SEM), with 95% confidence intervals reported for inferential statistics from LME model fits.

Prior to analysis, trials and participants were screened for data quality. Individual trials were excluded if the participant failed to initiate a movement, initiated a movement more than 200 ms prior to the go cue, failed to reach the target within 500 ms, or if the hand angle deviated by more than ±30° from the fixed target direction. Hand angle was defined as the angular deviation between the start–to–target vector and the movement direction measured at peak reach velocity. Participants were excluded if more than 33% of their trials met these criteria or if their behavioral pattern suggested a clear lack of instruction coherence. This resulted in the removal of two participants from the internal-monitoring experiment, four from the looming-cue experiment, and one participant from the numeric-cueing experiment. Overall, 12.01% of trials from the internal-monitoring experiment, 4.36% of trials from the looming-cue experiment, and 8.05% of trials from the numeric-cueing experiment were removed prior to the main analyses.

To quantify context-dependent learning while accounting for trial-by-trial adaptation, we fit linear mixed-effects models to baseline-subtracted hand angles during the training phase, separately for each experiment. For each participant, hand angles were baseline-corrected by subtracting the corresponding mean hand angle from the baseline phase separately for the 500 and 1500 ms conditions. Fixed-effects predictors included current-trial context, rotation direction on the preceding trial (n-1), rotation direction two trials back (n-2), and trial number, together with all interactions among the current context, previous-trial rotation, and trial number, plus the interaction between two-trials-back rotation and trial number. Trial number was mean-centered within participants but retained in units of trials. The model included participant-specific random intercepts and random slopes corresponding to the fixed-effects structure and was fit using restricted maximum likelihood. For visualization of context-dependent learning over training, we additionally regressed baseline-subtracted hand angle on the previous trial’s rotation direction separately for each participant and used the resulting residuals to compute the binned context effect between the 500 and 1500 ms conditions.

For the computational model analyses, goodness of fit for model predictions was quantified by the coefficient of determination (*R*^2^), computed from least-squares errors between model predictions and empirical data across all five test delays, and models were compared in their ability to predict the generalization in the test phase. Models were also compared at the participant-level using Wilcoxon signed-rank tests on each participant’s root mean squared error (RMSE) at the three untrained delays in the test phase. Model-fitting details, parameter constraints, and model implementation procedures are described in the next section.

### Computational Modeling

We compared two models that differed primarily in how temporal context was represented. Both models contained a delay-agnostic general motor adaptation component and a temporally specific learning component. Models were fit only to the training-phase data, and fitted models were then used to generate predictions for the no-feedback test phase.

#### Subjective timing model

This model combines two computational concepts: hazard rates (for temporal anticipation) and context-dependent associative motor adaptation (for motor learning).

For the timing component of the model, we assume that participants, upon seeing the ‘set’ cue, actively anticipate the next event – the ‘go’ cue. The go-cue event can occur at one of a set (*N*) of learned delay intervals, *μ*_*i*_, with subjective probability density over event time (*t*) modeled as a mixture of Gaussians centered on each interval,

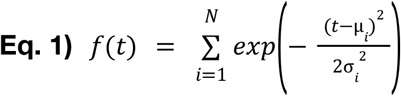

where widths are scaled with elapsed time via the Weber fraction *ω* (scalar timing), and *σ*_*i*_ = *ωμ*_*i*_. We assume uniform mixture weights, yielding a multimodal predictive distribution once the intervals are learned. We also assume that the specific intervals are learned rapidly during the baseline phase of the task.

The model converts this subjective probability density into a subjective hazard function *h*(*t*),

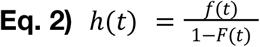

where *F* is the cumulative distribution function of the subjective PDF. Reaction time (RT) is then computed via the inverse of the hazard rate, with an RT floor at *a* and a linear scaling parameter *b*,

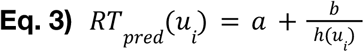

where *a* represents an RT floor and *b* controls the scaling between hazard and RT. Thus, a higher hazard rate (i.e., at later intervals) yields a shorter RT.

We modeled context-sensitive motor learning using a state space model of motor adaptation (Smith et al., 2006). The model contains a delay-agnostic generic state, *x*, that is active on every trial, and two delay-specific states, V_500_ and V_1500_, one for each trained set-go interval. On trial *n*, the model’s predicted hand angle is the sum of the generic state and whichever context specific state is currently active:

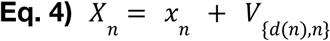

where *d(n)* denotes the delay condition on trial n. The generic state and the active context state are updated according to the difference between the fixed rotation magnitude, R (set to either ±15° to match the rotation used in the task), and the model’s predicted hand angle, *X*_*n*_ :

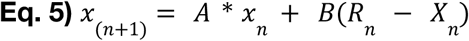

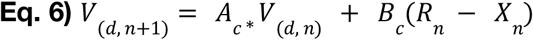

where *d* indexes the two discrete context states. Here, *A* and *A*_*c*_ are retention parameters and *B* and *B*_*c*_ are learning-rate parameters for the generic and context-specific states, respectively.

The delay-agnostic generic state essentially captures the near-catastrophic interference of having interleaved rotation signs, while the delay-specific states capture the incremental learning of temporal contexts for different motor memories (i.e., context effects). The two context states do not interact directly and share a single retention/learning rate pair, *A*_*c*_ /*B*_*c*_.

The timing parameters *ω, a* and *b* were fit to participants’ training-phase reaction times. Motor learning parameters *A, B, A*_*c*_ and *B*_*c*_ were fit to participant hand angle data in the training phase of the task using the *fmincon* function in MATLAB, minimizing the root-mean-squared-error between subjects’ actual hand angles during training and modeled hand angles. The lower and upper boundaries constraining each parameter during fitting were [0.05, 0.30] for *ω*, [0.10, 0.50] for RT floor (*a*), and [0, 1] for RT scale (*b*) in the timing model; and [0, 1] for *A*, [0, 0.5] for *B*, [0, 1] for *A*_*c*_, and [0, 1] for *B*_*c*_ in the state space model. The timing component of the model fit was repeated from 20 random initializations, and the motor-learning model fit from 100 random initializations, to reduce the likelihood of convergence on local minima. In the externally-cued experiment, reaction times did not differ across the two delays and hence carried no signal from which to estimate temporal uncertainty; *ω* was consequently fixed at 0.001, while all other parameters were fit as described above.

Finally, our primary goal was to estimate participant behavior in the testing phase. To that end, we conjoined the two model components, both of which were only fit to the training phase. We treated the asymptotically learned predictive value of each context state, *V*_500_ and *V*_1500_, as weights, which were then multiplied through the subjective timing PDF at the appropriate intervals. This resulted in a model that takes the subjective timing PDF (i.e., the Gaussian mixture with Weber scaling) and integrates it with memories of the upcoming temporal event’s likely ‘consequence’; that is, it estimates a continuous prediction of the ‘expected state’ (rotation) at each temporal interval as determined by the error-based learning model:

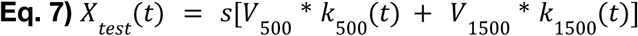

where *k*_500_ and *k*_1500_ are the Gaussian generalization kernels centered on 500 and 1500 ms and s is a fixed scale factor accounting for reduced magnitude of learning in the test phase (see below).

To illustrate: Where the subjective PDF has a value of zero, the predicted rotation is zero; where the subjective PDF has a nonzero value, a nonzero rotation prediction is made, whose sign is consistent with the sign of the learned motor memory *X* near that time point, and whose magnitude is proportional to the temporal separation between a probed interval and the trained intervals. Because contextual learning is gradual, the strength of the rotation predictions incrementally increases over training. The model predicts varying degrees of generalization behavior at unseen delay intervals, and the exact shape of that generalization function is influenced by both learning dynamics and Weber scaling. When reaction times do not differ meaningfully between the long and short delay, as seen in the externally-cued condition where participants infer the context from the start of each trial, the model is fit with a Weber fraction near zero. Since the width of the Gaussian component in the subjective PDF scales with this parameter (*σ*_*i*_ = *ωμ*_*i*_), a near-zero Weber fraction causes the PDF to collapse into two narrow, non-overlapping peaks centered on the trained intervals. The model therefore predicts negligible generalization to untrained delays under these conditions. Lastly, a single additional scale factor (s = 0.5) uniformly shifted the testing phase prediction into an appropriate scale, capturing forgetting and memory decay.

#### Granule-cell basis model

Inspired by recent cerebellar models of interval timing (e.g., Narain et al. 2018), we developed a second model in which elapsed time was continuously represented by a distributed population of overlapping, granule-cell-like temporal basis functions. Because nearby time points activate overlapping populations of these basis functions, learning associated with one delay will naturally generalize to neighboring delays.

This model shares the same overall architecture as the subjective timing model described above: a delay-agnostic generic state combined with a delay-specific learned component, updated by the same error-driven state space rule. The major difference lies in how context is represented. The two discrete context states, *V*_500_ and *V*_1500_, are replaced by a continuous population of temporal basis functions, ***ϕ***_j_ (t), with preferred time *c*_j_ distributed from 250 to 1750 ms in 100 ms increments. Rather than a single scalar value per context, learning now updates a full weight vector, w, spanning this basis population.

Following the general temporal-basis framework of Narain et al. (2018), different basis functions were tuned to different elapsed times, providing a distributed representation of the preparatory interval. The parameters governing this temporal basis set were fixed across participants and experiments and were chosen to reproduce the broad qualitative properties of the TRACE model (Narain et al., 2018); namely, broader and lower amplitude tuning at longer intervals. The activity of each basis function *j* at elapsed time *t* was defined as:

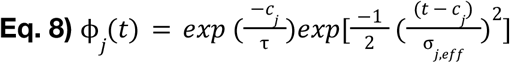

where *τ* controls the gradual decay in amplitude of the basis functions tuned to later times. We fixed *τ* at 10,000 ms. The width of the basis functions increases with their preferred intervals according to:

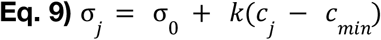

where σ_0_ = 100 ms represents the baseline width of the temporal basis functions and k = 0.35 determines the increase in tuning widths at later times. We also incorporated a fixed timing noise term, σ_*noise*_ = 150 ms, into the effective width of each basis function:

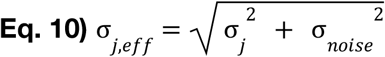

On trial n, the model’s predicted hand angle was given by the sum of the delay-agnostic component and the weighted activity of the temporal basis functions at that trial’s experienced delay interval:

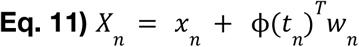

where *X*_*n*_ is the model-predicted motor output on trial n, *x*_*n*_ is the delay-agnostic state, and 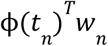 is the temporally specific contribution from the basis functions. The delay-agnostic state and the temporal weights were updated according to the difference between the signed rotation magnitude, *R*_*n*_, and the model output:

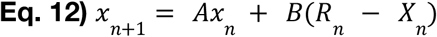

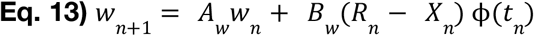

Here, *A* and *A*_*w*_ are standard retention parameters and *B* and *B*_*w*_ are standard learning rate parameters for the delay-agnostic and temporally-specific components, respectively.

The four free parameters (*A, A*_*W*_, *B* and *B*_*W*_) were fit separately to each participant’s training-phase hand angle data using MATLAB’s *fmincon*. Parameter bounds were [0,1] for *A*, [0,0.5] for *B*, [0,1] for *A*_*W*_ and [0, 0.1] for *B*_*W*_. Fits were repeated from 100 random initializations, and the parameter set yielding the lowest training-phase RMSE was retained.

At the end of the training, the delay-agnostic state and temporal basis weights were held fixed. Test-phase predictions were generated by evaluating the temporal basis population at each tested delay:

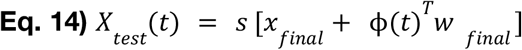

Thus, generalization to untrained delays emerged from overlap between the temporal basis functions activated at both the trained and untrained delays. No additional parameters were fit during the test phase. Identical to the subjective timing model, test-phase predictions were multiplied by the same scale factor (*s* = 0.5) to account for the reduced magnitude of effects during the test phase. Estimating *s* from the trained delays instead yielded values near 0.5 in both experiments (0.48-0.58) and did not change the model comparison.

## Acknowledgments

A.S., H.H., and S.D.M. designed the experiments. A.S. performed the experiments. A.S. and S.D.M. performed analyses and modeling. A.S. and S.D.M. drafted the manuscript, and all authors edited it. We thank the ACT Lab for helpful discussions. Work supported by grant R01 NS134754 (S.D.M.) from the National Institutes of Health.

## Notes

### Competing Interest Statement

The authors have declared no competing interest.

