## Supplementary Material for "Temporal Indexing of Motor Memory"

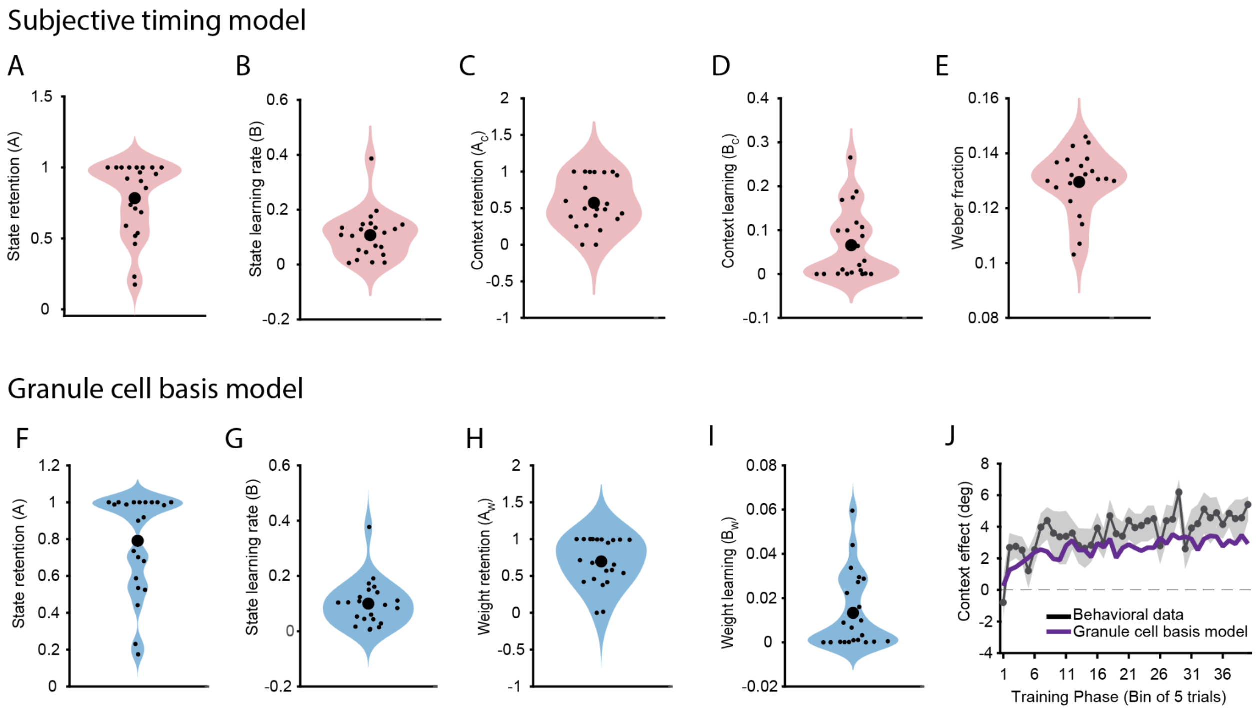


**Figure S1**: Fitted model parameters and learning fit for internally-monitored task. **(A-E)** Fitted parameters for subjective timing model, comprising the timing component and error-based learning component. **(A)** State retention, **(B)** State learning rate, **(C)** Context retention, **(D)** Context learning rate, **(E)** Weber fraction, estimated from participants' training phase reaction times. **(F-I)** Fitted parameters for granule-cell basis model. **(F)** State retention, **(G)** State learning rate, **(H)** Weight retention, **(I)** Weight learning rate. **(J)** Binned context effect across training phase comparing behavioral data (mean ± SEM) to the granule-cell basis model's simulated hand angle. Individual dots show participants' fitted parameter values and larger dots show the group mean. We note that both the subjective timing and granule-cell basis models share the same underlying error-based learning model structure, and differ only in their temporal representation and test phase predictions (see *Methods*).

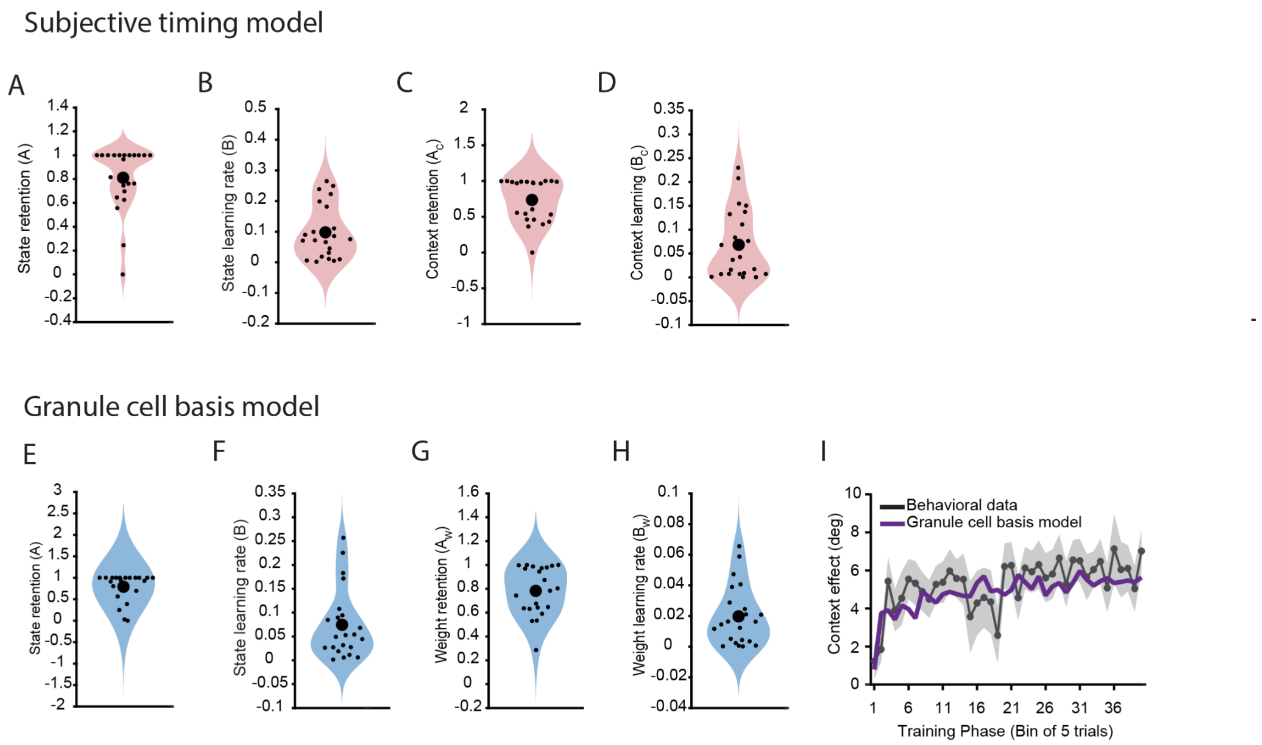


**Figure S2**: Fitted model parameters and learning fit for externally-cued task. **(A-D)** Fitted parameters for subjective timing model, **(A)** State retention, **(B)** State learning rate, **(C)** Context retention, **(D)** Context learning rate. **(E-H)** Fitted granule-cell basis model parameters, **(E)** State retention, **(F)** State learning rate, **(G)** Weight retention, **(H)** Weight learning rate. **(I)** Binned context effect across training phase comparing behavioral data (mean ± SEM) to the model's simulated hand angle. Individual dots show participants' fitted parameter values and larger dots show the group mean.

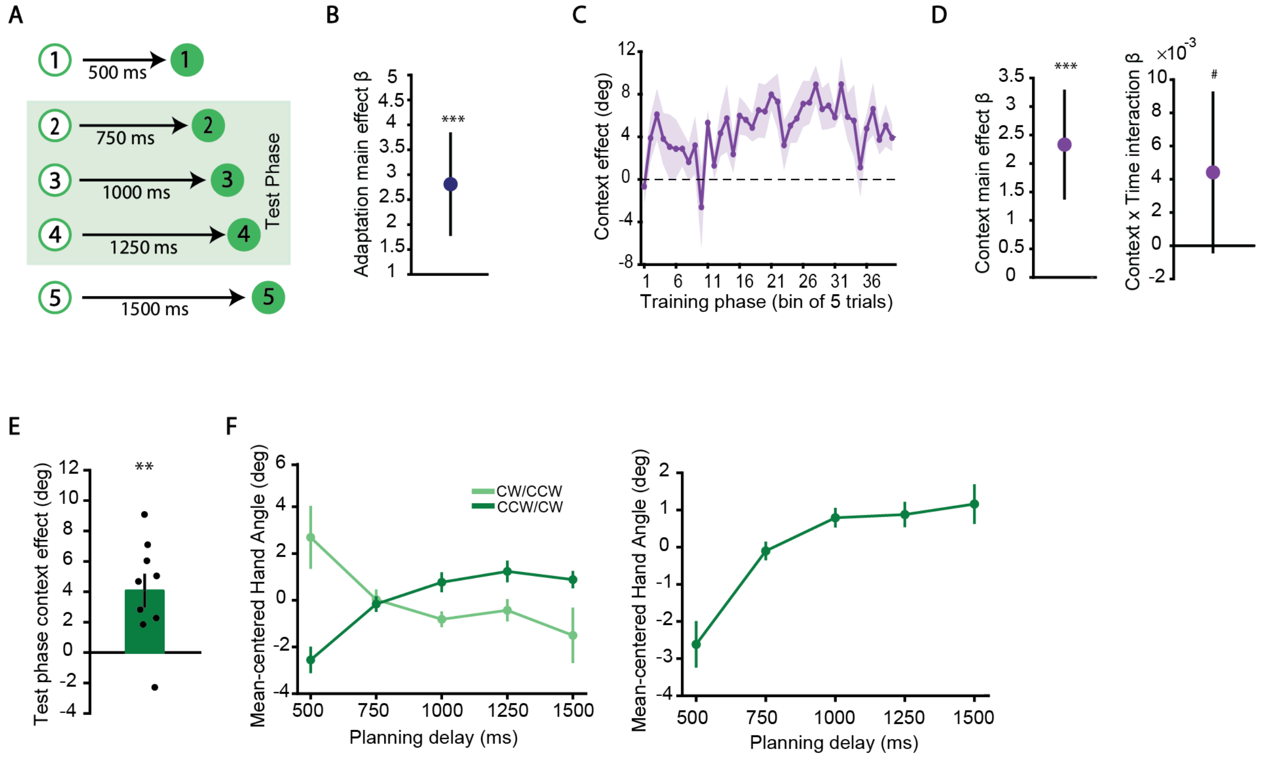


**Figure S3**: *Results: Discrete numeric cueing experiment.* (**A**) Schematic of the discrete temporal cueing task. Numbers explicitly indicate the upcoming planning delay, with intermediate values corresponding to the novel delays during the test-phase. (**B**) Trial-by-trial adaptation during training, shown as adaptation main-effect β coefficients. (**C**) Temporal context effects across training phase after accounting for adaptation, plotted in bins of five trials. (**D**) Regression coefficient characterizing contextual learning during training. Left: context main-effect β. Right: context X trial # interaction β. (**E**) Context dependent hand angle separation in test phase. Bar graph depicts the group mean context effect, with individual points showing each participant's effect. (**F**) Left panel: mean-centered hand angle across the five tested delays for the two counterbalanced subgroups plotted separately in the test phase; Right panel: same test-phase data after aligning the subgroups to a common delay-rotation mapping. Error bars = 1 S.E.M. # marginal effect, $^{**}p < .01$, $^{***}p < .001$
